# Movi Color: fast and accurate long-read classification with the move structure

**DOI:** 10.1101/2025.05.22.655637

**Authors:** Steven Tan, Sina Majidian, Ben Langmead, Mohsen Zakeri

## Abstract

The number of reference genomes is rapidly increasing, thanks to advances in long-read sequencing and assembly. While these collections can improve the sensitivity and specificity of classification methods, this requires highly efficient compressed indexes. K-mer-based approaches like Kraken 2 are efficient but limit the analysis to a fixed k-mer length. This is hard for the user to set ahead of time, and suboptimal settings can harm sensitivity and specificity. Methods that use compressed full-text indexes like SPUMONI2 and Cliffy lift this constraint, but are less efficient than k-mer-based tools. Further, these methods either cannot report a full listing of genomes where a match occurs, or cannot scale to large reference databases.

We propose new methods and algorithms that use compressed full-text indexes to enable multi-class and taxonomic classification. Unlike past compressed-indexing methods for classification, ours uses the move structure, which is extremely fast thanks to its locality of reference. Our method, called Movi Color, augments the main table of the Movi index. Specifically, Movi Color assigns a “color” to each run of the Burrows-Wheeler Transform according to the subset of genomes from which the run suffixes originated. When the reference is highly repetitive – as is typical when indexing pangenomes or reference databases – only certain colors occur, creating opportunities to compress the index. For species-level classification, Movi Color achieves over 1.6*×* higher precision and 2*×* higher recall than Kraken 2 and Metabuli. At the genus level, it achieves 70% higher precision and 80% higher recall. Movi Color’s read processing time is 7-20× faster than Metabuli and is a comparable to Kraken 2. Although Movi Color uses more memory than both Kraken 2 and Metabuli, its speed-accuracy trade-off makes it well-suited for real-time or high-throughput scenarios.

## 1 Introduction

Taxonomic classification aims to identify the likely origin of each sequencing read in a complex metagenomics sample [9, 1]. Numerous methods have been developed to achieve accurate and efficient methods for this task. Efficiency is increasingly a prime concern as advances in long-read sequencing have enabled the rapid creation of new reference genomes and pangenomes. Most methods first create an index over the reference sequences from a wide range of taxonomic groups. Sequencing reads can then be queried against the index to determine the best taxon match. Methods like Kraken [31, 30] and Clark [25] use k-mer-based indexes, which store all substrings of length k (k-mers) from the reference sequences, along with information about the taxonomic groups in which each k-mer appears.

An advantage of k-mer-based indexes is that their size grows primarily with the number of distinct k-mers in the collection, i.e. the amount of non-redundant sequence. As new strains of the same species are added to the index, the size grows only by the number of k-mers that span sites with not-yet-seen variation, rather than by the total number of k-mers. These methods are also very fast, since the k-mers, or hash values derived from k-mers, can be used as keys for an associative data structures like hash tables. A limitation, however, is that choosing a single k value is often insufficient to capture both short conserved regions and long discriminative matches, especially when classification must operate across varying evolutionary distances [17]. For instance, as the size of reference databases grows, k-mer-based classifications become less specific [21].

Recently, compressed full-text indexes like the *r*-index [20] have been used for taxonomic classification. At *r*-index’s core is a run-length-encoded representation of the Burrows Wheeler Transform (BWT). For repetitive inputs, the number of BWT runs increases more slowly than the length of the input, similarly to how the number of distinct k-mers also grows more slowly [32]. Because *r*-index is a full-text index, it does not depend on a single k-mer length; rather, it can find matches of any length. Past work shows that this leads to classification accuracy consistently higher than that of k-mer-based approaches [4, 3, 29].

For taxonomic classification, SPUMONI2 stores genome-of-origin information for the suffixes on BWT run boundaries. This allows it to tally partial classification information over the course of the pattern-matching process. Cliffy uses a run-compressed representation of the longest common prefixes (LCPs) of the suffixes with every possible genome of origin. Both methods can be combined with minimizer digestion to improve query speed and reduce index size. However, while they achieve higher classification accuracy compared to Kraken 2, their query speed remains significantly slower, and their indexes are much larger. Additionally, SPUMONI2 cannot provide a complete listing of all the genomes where a read matches. Further, Cliffy cannot list all the genomes where a match occurs, but can accurately report the lowest common ancestor (LCA) of those genomes within a taxonomic tree.

Full-text indexes can be used to compute matching statistics (MSs) for a read with respect to an index, where MSs include the length of longest exact matches between all suffixes of the read and reference sequences [28]. Ahmed et. al proposed pseudo-matching lengths (PML) as an approximation of MSs, which are faster to compute and can be used for accurate binary classification [2]. Zakeri et. al [32] showed that PML computation can be made drastically faster by using a move-structure index [23].

Here we extend Movi to include “run colors,” encoding the origin of the suffixes in each run. This is inspired by how color information is used in colored de-bruijn graphs [15, 19]. In that setting, a color denotes the particular subset of genomes or taxa in which a k-mer occurs. Also inspired by past work on colored de-bruijn graph, we attempt to compress the color information using the fact that some colors are frequently re-used across BWT runs.

We hypothesize that run colors encode powerful, compressed information that can be sufficient for accurate multi-class classification. To test this, we apply the Movi Color classification scheme to the task of taxonomic classification of long reads. Our results show that Movi Color achieves higher classification accuracy than Kraken 2 while being almost as fast. Although Movi Color requires significantly more memorye compared to Kraken 2, we note that there is substantial potential for further compression—both in the color representation and in the underlying Movi index.

## 2 Methods

### 2.1 Background

#### 2.1.1 Burrows-Wheeler indexes

Let S be a string of length |*S*| = *n* over the ordered alphabet Σ + {$}. Assume *S*[*n* − 1] = $ is a unique terminator character that is lexicographically smaller than the others. A suffix of S starting at position *i* is denoted by *S*[*i*..*n* − 1]. The Burrows-Wheeler Transform (BWT) is a permutation of *S*’s characters according to the lexicographical order of the suffix of S starting from the character’s right. For example, *S*[*i*] comes before *S*[*j*] in BWT, if and only if *S*[*i* + 1..*n* − 1] is lexicographically smaller than *S*[*j* + 1..*n* − 1] [11]. Figure 1.a illustrates the BWT of a string created by concatenating 3 other strings. We use the term “document” to describe a sequence or group of sequences that constitutes the most specific classes we can use in our classification strategy. In our example, each input string is a separate document, which we call *D*_1_, *D*_2_, and *D*_3_. Figure 1.b shows how the characters in the BWT are sorted based on the suffixes in the original string S.

**Figure 1.**
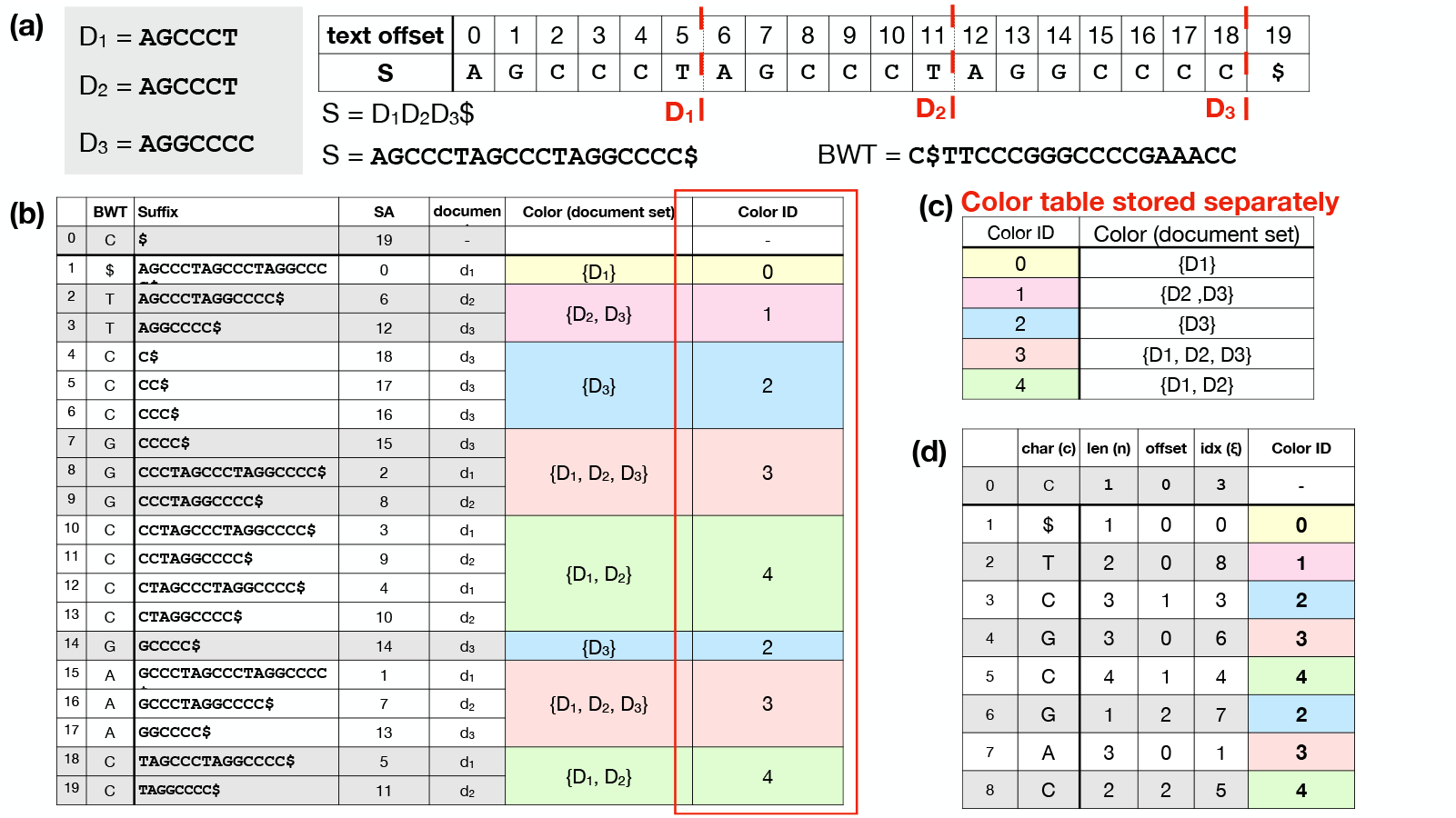
(a) The documents and the concatenated text used for building the BWM. The red dotted lines are the document boundaries in the concatenated text. (b) The BWT entries with the corresponding suffixes and suffix array (SA) entries. The document ID at which the suffix starts is assigned based on the SA entry and the document boundaries in the concatenated text. The color of a run is the set of all the documents from the BWT offsets in the run. Each separate color is assigned a unique ID. The color ID column (indicated by the red box) is stored in the Movi index as a new column in the main table. (c) The color table which maps a color ID to a set of documents representing the color is stored separately in the index. (d) The Movi Color table which is the same as the Movi table with an additional column to store the color ID for each run.

The Burrows-Wheeler Matrix of S, which we call BWM, is a matrix whose rows are S’s distinct cyclic rotations in lexicographical order. The last column of the BWM is the BWT. The BWM’s first and last columns are related by the Last-to-First mapping (“LF-mapping “), which states that the *i*^th^ occurrence of a character c in the last column of the BWM corresponds to the same text occurrence as the *i*^th^ occurrence of c in the first column. The FM-index uses the BWT plus some auxiliary structures to perform LF-mapping and match queries efficiently.

The BWT permutes *S* so that the characters with the most similar right contexts appear consecutively. The BWT of a repetitive input will tend to have long same-character “runs” [20]. The number of maximal equal-letter runs is denoted *r*. The *r*-index is a compressed full-text index based on the run-length-encoded BWT. Variants of the *r*-index achieve a space complexity of *O*(*r*) machine words, or a time complexity of *O*(*m*) queries, where *m* denotes the query length |*R*| [14]. While these are favorable bounds, the *r*-index can be slow in practice due to poor locality of reference. That is, an individual index query requires accesses to many data structures such as bitvectors, wavelet trees (and their constituent bitvectors), and indexes enabling efficient rank and select queries over the bitvectors. This leads to many cache misses and a slower query in practice.

The move structure is an alternative compressed full-text indexing structure. It offers *O*(*r*) space and constant-time matching queries (i.e. LF-mapping queries) [23]. Movi constructs an index based on the move structure and tailored to DNA pangenomes [32]. The index consists of a table (*M*) with each row corresponding to a BWT run^2^. Each BWT offset is represented by a pair (*i, j*), denoting an offset *j* into a BWT run index *i*[10]. Each row of the Movi index encodes the character of the run *c*, the run length *n*, and a tuple (*ξ*, offset) denoting the LF-mapping destination of the run head. Additionally, Movi can store a data structure (the thresholds data structure) that allows for some specific LCP queries [7], which are in turn used in the computation of MSs and PMLs. An example of the Movi table is illustrated in Figure 1.d.

#### 2.1.2 Index queries for matching and classification

Let *R* = *R*[0..*m* − 1] be a query string (e.g. a DNA sequencing read) of length *m*. The algorithm processes the letters of the query right-to-left, starting with *R*[*m* − 1]. At step *i*, the algorithm computes which interval of the BWM consists of rotations having *R*[*m* − 1 − *i*..*m* − 1] as a prefix. If the interval becomes empty at step *j*, that indicates that the suffix *R*[*m* − 1 − *j*..*m* −1] does not occur in S. Figure 2 illustrates the process.

**Figure 2.**
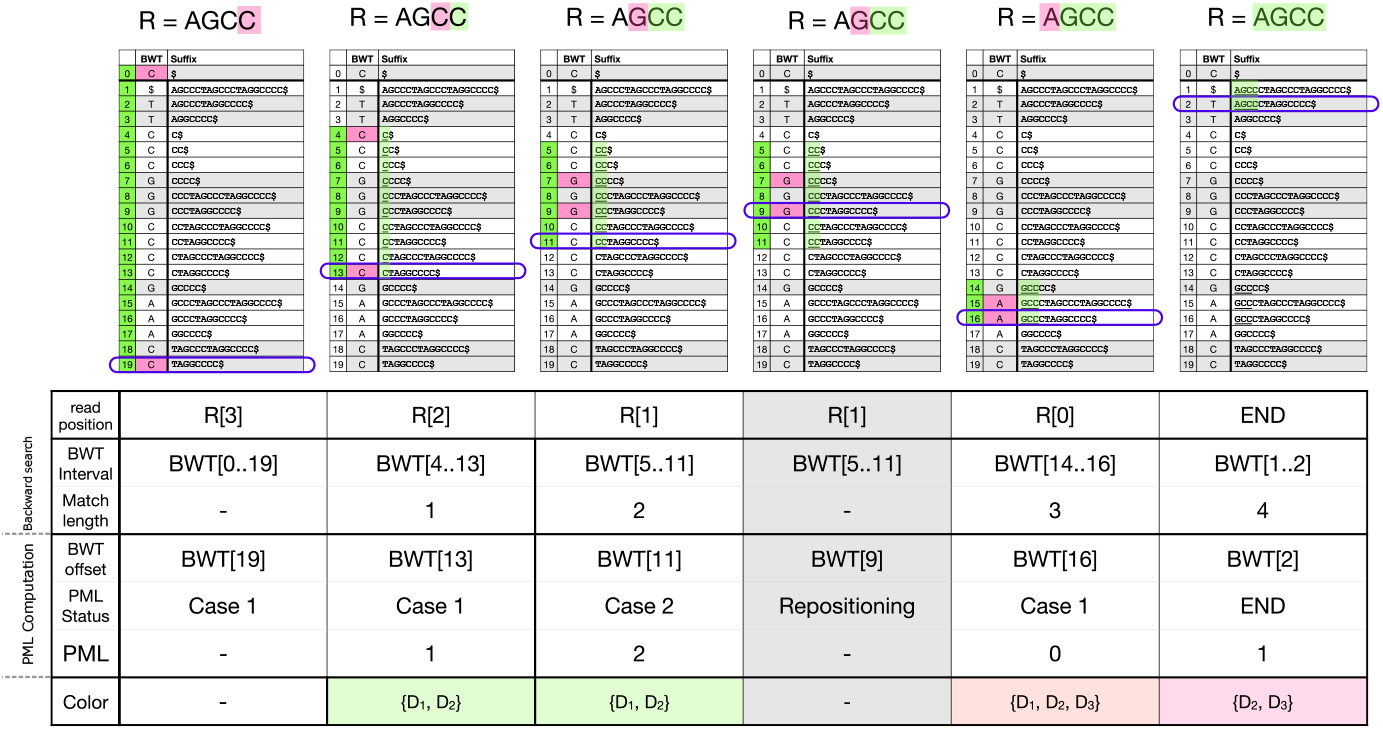
An example to compare backward-search and PML computation algorithms side by side. The read *R* = “*AGCC*” is being searched to the index starting at the right most end. The backward search maintains a BWT interval (BWT[i..j]) which contains all the possible exact matches. The offsets in the BWT interval are showed by green at each steps. The shaded green characters are the ones that are matched by the backward search until that step. The pink character is the character queried at each step. The purple box shows the offset tracked by the PML query at each step. When the character of the BWT offset matches the read’s character, we have case 1, otherwise it is called a case 2. The PML length is incremented by one for each case 1 and is reset to 0 by case 2. The repositioning is performed after each case 2 to find a BWT offset that matches the read’s character. The bottom row of the table shows the colors identified by the tracked BWT offset during PML query. See Figure 1 for the definition of colors.

When the query is a long read, backward search might match only a short suffix of the read before reaching a suffix that does not occur. If the goal is to classify the read, we could continue the process by initiating a new backward search round starting from the offset that caused the interval to become empty in the previous round. Repeating this process until we reach the left-hand side of the read will yield a sequence of half-maximal exact matches (half-MEMs) covering the R. Taking the idea a step further: we could initiate a separate backward search process from *every* read offset, allowing us to find all half-MEMs, from which we can in turn infer the maximal exact matches (MEMs). We call this “all-offsets backward search.” This would be an inefficient strategy, requiring a number of matching steps equal to the total length of all half-MEMs.

MONI [28] proposed an efficient *r*-index-based algorithm for finding matching statistics (MSs), i.e. the half-MEMs found by all-offsets backward search. Later, pseudo-matching lengths (PML) were proposed as an alternative to MSs that are easier to compute [2]. The PML-finding algorithm processes the read from right to left, maintaining a specific BWT offset (rather than an interval) at each step. Initially, the BWT offset is set arbitrarily; by convention, we set it to the final element of the BWT, BWT[n-1]. At each step, the character of the current BWT offset either matches the next character in the read (case 1), or does not (case 2). In case 1, the BWT offset is updated using the LF-mapping. In case 2, a repositioning step is conducted before performing the LF-mapping. Repositioning examines nearby rows (either up or down) to find a BWT offset that matches the read’s character. The thresholds in a BWT run, determine for each BWT offset within the run whether up or down repositioning reaches an offset with a higher LCP. Pseudocode is given in Algorithm 1. Figure 2 illustrates the BWT offset tracked during the algorithm, and how it relates to the larger interval that would have been computed with all-offsets backward search.

### 2.2 Adding colors

Suppose we are indexing documents *D* = {*D*_1_, *D*_2_, …}, as in Figure 1.a, with the goal of classifying reads according to their likely document(s) of origin. To accomplish this, we wish to augment the Movi data structure with information about document of origin. A naive approach is to store a document ID for each offset of the BWT, but this would grow linearly with the input length, which could be expensive in practice. Instead, we store one “color” per run, where each color is a distinct subset of the documents. Consider the *i*-th BWT run, which corresponds to consecutive rows *j*_1_, …, *j*_*k*_ in the BWM. We define run *i*’s color *C*_*i*_ to be {*D*_*x*_ : *x* ∈ {*j*_1_, …, *j*_*k*_}}, the set of all document IDs for its rows. Note that *C*_*i*_ is a set; we do not encode how many times a document occurs in the run.

#### Algorithm 1

Pseudo-matching length (PML) computation for read R using Movi table M. A row of M encodes a character (c), a length (n), and a threshold for every alphabet character. An offset is represented by a pair (i, j) denoting offset j into the run with index i. The RepositionUp and RepositionDown functions scan nearby rows of M’s to find the nearest row having character c.

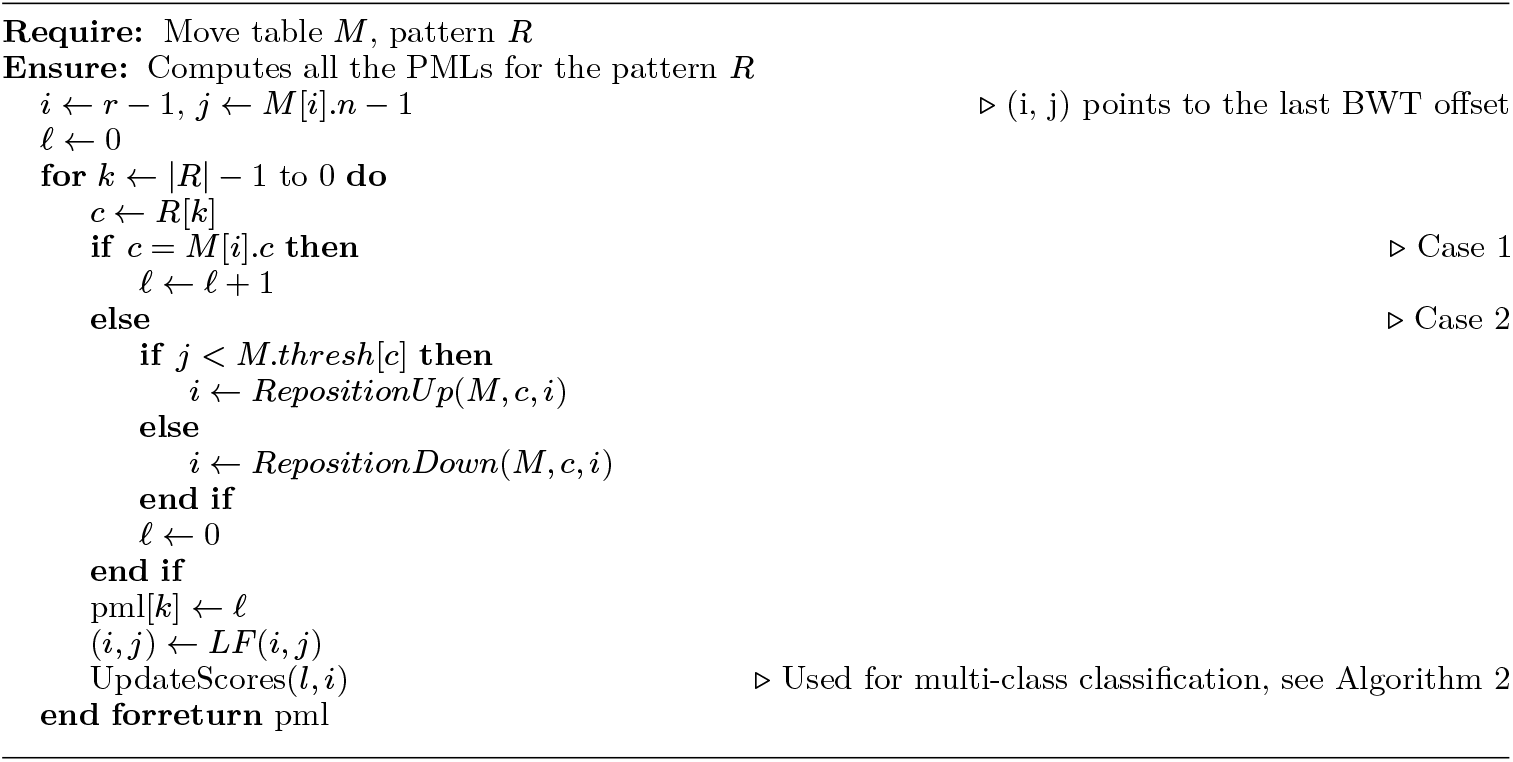

The definition of the run colors is motivated by the colored de-bruijn graph, where a k-mer’s color similarly represents the set of documents where it appears [15]. Figure 1.b illustrates the per-run colors in the context of the BWM. Note that multiple runs can share the same color. For example, the colors for the 4th (“CCC”) and 7th (“G”) runs are both {*D*_3_}. To avoid redundancy, we assign a unique color ID to each distinct color and store only the color ID for each run. We define *C* = {*C*_1_, *C*_2_, …} as the set of all the unique colors that exist in the index. Figure 1.c shows the “Color Table” which maps color IDs to the corresponding document sets, and Figure 1.d shows the Movi Color table, which includes an additional column to record the color ID associated with each row.

We define the shared-LCP of a run as the length of the longest prefix shared by all the rows within a BWM run. We expect that shared-LCP of a run is typically long; specifically, it should usually be longer than the length of a random match between a query and the reference collection. By associating a color with a run, we can confidently link similar sequences (the shared-LCP of the run) to a set of documents. In this sense, the shared prefixes of a run are analogous to a k-mer, but with the distinction that the length of the shared-LCP varies based on how similar the rows within the run are. Biologically, we may imagine that conserved regions among genomes tend to cluster in runs, and thus the genomes that share conserved regions may commonly appear together in the same Movi Color.

#### 2.2.1 The color table representation

The color table is a lookup table that associates colors with the set of documents represented by the color. The size of the table depends on the number of distinct colors, |*C*|, and the representation used for each color. A straightforward representation would be to use a bitvector of length |*D* |, with each bit indicating the presence or absence of a document. In that case, the table occupies |*C*| × |*D*| bits. Since the number of colors never exceeds the number of runs, this space for this representation is also no more than *r* × |*D*| bits. However, when the documents are diverse (e.g. each document represents a distinct species) and |*D*| is large, the color bitvectors contain mostly 0’s. In that case, it can be more efficient to store offsets of the set bits (a sparse representation) rather than the full bitvector (a dense representation) [13]. If *b* denotes the average number of set bits in all the color bitvectors, the total space then becomes |*C*| × *b* integers.

For repetitive inputs like pangenomes, we expect that many runs will share the same color. This has been observed in colored de-bruijn graphs [6, 13]. The ratio of the number of runs to the number of distinct colors, *r/* |*C*| is proportional to the benefit achieved by using the color tables instead of storing the colors explicitly for each run.

### 2.3 Multi-class classification by PMLs and colors

We propose a method that augments the usual PML-finding algorithm to additionally detect and tally document information. Specifically, our proposed algorithm will tally the colors of the runs that the PML process moves through and, for each color encountered, increase a score associated with the documents that comprise the color. Once PML finding is done, the document with the highest score is selected as the most likely origin of the read. Alternately, the algorithm can report all documents having scores close to the highest score, and we can classify the read as having originated from the lowest common ancestor of those documents, enabling taxonomic classification at higher levels.

For a length-m query, the PML algorithm yields a sequence of lengths *L*_0_, *L*_1_, ..*L*_*m−*1_ such that *L*_*k*_ indicates there is a match of length at least *L*_*k*_ between *R*[*k*..*m* −1] and a row in the BWM. When the length *L*_*k*_ is greater than 0, Movi Color looks up the color of the current BWT run. This pseudo code of the update strategy is provided in Algorithm 2. As an example, the lengths and their associated colors are shown in the bottom two rows of table in Figure 2. Each time we encounter a color, we increment a score for each of the color’s included documents. At the end of the example shown in Figure 2, the final scores would be: *D*_1_ : 2, *D*_2_ : 3, and *D*_3_ : 1. Note that the color associated with BWT[16] is not considered for incrementing the scores, because the associated PML is zero.

This document scoring strategy can make two categories of errors, both flowing from the fact that the PML algorithm tracks a single row in the BWT, whereas the colors describe entire runs.

While this strategy does not guarantee scoring all documents that match the query, incrementing the scores of the documents present in the run color of the tracked row increases the chance of capturing additional true matches, if they exist. As an example in Figure 2, *R*[2..3] = “*CC*” matches all three documents as the backward search range suggests. But the tracked PML row (BWT[11]) only indicates that *D*_2_ matches R[2..3]. When we consider the color of the run including BWT[11], *D*_1_ is also considered as a possible origin of the match. However, we are still missing *D*_3_ which also includes the match.

On the other hand, this approach may increment the score of a document that does not contain the match. That happens when the shared-LCP of the run is shorter than the length of the match. In Figure 2 the color of the final run is {*D*_2_, *D*_3_}, therefore the score of both documents are incremented but only *D*_2_ includes the observed match.

We argue that the second type of error is very unlikely to be problematic in practice, particularly because we ultimately consider only the documents with the highest scores as candidates for classification. Importantly, the document associated with the tracked BWT row is always included in the color. Excluding highly pathological cases, when another document appears in the colors as frequently as the true document, it is likely that this document is a valid match for the read; otherwise, we would expect it to occur in only a fraction of the observed colors along side the true document. In Figure 2, document *D*_1_ appears in 2 out 3 colors considered during PML matching and is indeed a true match for the read. In contrast, document *D*_3_ appears in only one of the colors, as it does not contain the true match.

#### 2.3.1 Repositioning without thresholds

Regular PML computation relies on the thresholds structure to determine the repositioning strategy when the character in the BWT offset does not match the read (case 2). The purpose of repositioning is to move to a nearby row in the BWM that has the highest LCP with the current row, thereby preserving the longest possible match.

While thresholds enable precise repositioning by guiding PMLs into regions of the BWM more closely related to the read, computing this structure may not always be feasible due to its cost. To address this, we explore two alternative strategies for repositioning in the absence of thresholds.

The first strategy is to always reposition to the up direction, effectively treating the threshold as if it were at the last position of the run. The second strategy places the threshold midway through the run, repositioning upward or downward depending on whether the BWT offset position is above or below this midpoint.

The intuition behind these strategies is that even if repositioning is not to the right direction, we still reach a row sharing a relatively long common prefix with the current match, because adjacent rows within BWM typically exhibit high LCP values with one another. Moreover, setting the thresholds to the middle of the run is reasonable heuristic. In particular, for long runs, rows near the beginning of a run are more likely to share longer prefixes with the rows above the run, while rows near the end tend to be more compatible with the rows below.

#### 2.3.2 Document listing and classification at higher levels

The documents used in the colors of the Movi Color index can be defined at any level of the taxonomy. For example, if the most specific classification we require is at the species level, all genomes in that species can be grouped into a single document, encoded by a single bit of the color bitvector.

Furthermore, when a read has more than one document with a significantly higher score than the rest, Movi Color can report all such documents. Specifically, Movi Color reports the document with the highest score, *D*_*best*_, along with any other document whose score is at least 95% of the score of *D*_*best*_. The taxonomic classification for that read is then assigned to the Lowest Common Ancestor (LCA) of all the reported documents in the taxonomy tree. Based on this strategy, Movi Color classifies the reads either at the level of the documents or at a higher taxonomic level. By default, Movi Color only considers *D*_*best*_, and at most one additional document that has the next best score greater than 95% of *D*_*best*_’s score. If no such second-best document exists, only *D*_*best*_ is reported. *D*_*best*_ will be the assigned classification for the read, which is at the lowest possible level defined by the documents.

##### Algorithm 2

Function for updating scores during multi-class classification (called from Algorithm 1). Takes the matching length and run index as input, and updates the scores of the documents in the run color.

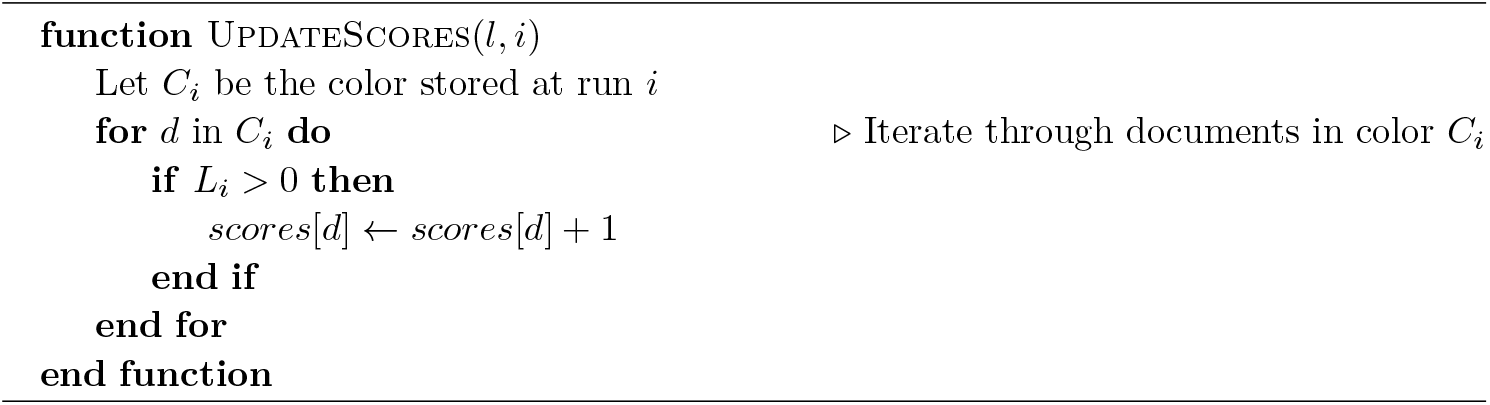

#### 2.3.3 Binary classification

We have developed a binary classification policy based on the PML values, to determine if a read is from the index before being considered for multi-class classification. In multi-class classification, this binary procedure helps exclude reads that is believed not to be in the index, avoiding unnecessary refined computations.

For each read, we compute the average PML length, which is the sum of all PML lengths divided by the length of the read. We classify a read as being from the index if the average PML is above a cutoff, and unclassified otherwise. To choose a suitable cutoff, we performed an empirical analysis which involved a set of null reads (which do not belong to the index) and a set of positive reads that match some documents in the index. For each set of reads, we compute the distribution of average PMLs. We then picked cutoff to be the 95-th percentile of the average PMLs from the null reads.This corresponds to a level-0.05 significance test, in which we allow 5% Type I Error (classifying a null read as part of the index). From the empirical analysis, this provided a power of over 98.5% (probability of classifying a read correctly to be part of the index).

## 3 Datasets and evaluation metrics

### 3.1 Genome datasets

We evaluated Movi Color using two reference datasets. The first we call the “48-species” dataset, which we generated, spanning 48 species within 3 kingdoms in the domain of Bacteria. This includes 31 species from *Pseudomonadati*, 16 from *Bacillati*, and 1 from *Fusobacteriati*. A full list of the species are available in the GitHub repository. The distribution of species across kingdoms roughly reflects the availability of reference genomes in NCBI. We downloaded 100 reference genomes for each of these 48 species, prioritizing complete assemblies where possible.

For each species, one randomly-selected genome was held out to be used for simulating reads, and the remaining 99 were included in the index, yielding a total of 99 × 49 = 4,851 genomes. We aimed to capture both redundancy (by including multiple genomes per species) and variability (by including diverse taxonomic groups). The total length of the genomes is 12.1 Gbases and the *n/r* ratio is 27.9.

The second dataset consists of all complete genomes available for species in the Phylum *Pseudomonadota*. This Phylum was selected because it accounts for approximately half of the bacterial genomes available in NCBI. The dataset includes all complete genomes from well-studied species *Escherichia coli* and *Salmonella enterica*, for example. We obtained the complete genomes of *Pseudomonadota* from a Kraken2 library which was compiled in November 2024 over all complete bacterial genomes in NCBI. The dataset includes a total of 75, 166 genomes from 5, 326 species. The total length of the reference sequences is 131 Gbases and the *n/r* is 10.4.

### 3.2 Read datasets

We evaluated the classifiers using two read datasets, one that we simulated and one from the Critical Assessment of Metagenome Interpretation (CAMI II) challenge [18]. The simulated dataset consists of ONT-like long reads simulated by PBSIM [24] using the R94 chemistry model. There are both positive and negative reads included in the simulated dataset. The positive reads are generated from all the species present in the 48-species dataset. The negative reads are simulated from all the species in *Thermotogati* kingdom which are absent from the index. There are a total 27, 158 long reads in the sample with an average length of 8, 939.

We also used the plant-associated metagenomic read dataset from the CAMI II challenge [18]. This sample includes 1, 691, 142 long reads with an average read length of 2, 956.

### 3.3 Evaluation criteria

To evaluate methods, we compared the estimated taxonomic classification with the ground truth. We followed the definitions of precision and recall used by tools such as Kraken2 [30]. Recall is defined as TP / (TP + VP + FN + FP), and precision is defined as TP / (TP + FP). When comparing at the species rank, a true positive (TP) is a correct classification at the species rank or lower. A vague positive (VP) refers to a correct classification at an ancestor of the true species when the classifier declines to report at a more specific rank. A classification is considered a false positive (FP) when it is incorrect at the species rank. A false negative (FN) occurs when a tool fails to report any taxonomic classification, even though the read originates from a species present in the index.

## 4 Results

### 4.1 Number of colors

We first analyzed the number of unique colors in Movi indexes built from different types of genome datasets. This experiment includes three datasets: the 48-species, the *Pseudomonadota*, and a dataset consisting of 50 genomes of *Bacillus subtilis*. In the first two datasets, introduced in Section 3.1, the documents are defined at the species level, meaning that all the genomes for that species are treated as one document together. For the *Bacillus subtilis*, each genome is treated as a separate document.

Table 1 shows the number of unique colors for each dataset. Notably, in all three cases, the number of colors is substantially smaller than the number of BWT runs. Specifically, the measure of *r/* |*C*|, the number of runs over the number of unique colors, is 56 for *Bacillus subtilis*, 760 for the 48-species, and 10 in the *Pseudomonadota* index. These numbers suggests that for different type of datasets, always many runs share the same color and using the color representation (adding only a color ID in a Movi row rather than the entire document set) is an efficient approach for storing the colors for all the runs.

**Table 1.** The number of colors and runs for the 3 indexes: 50 genomes from *Bacillus subtilis*, 48 diverse bacteria species with 100 genomes each, and complete genomes of *Pseudmonadota*.

| Dataset | *Bacillus subtilis* | 48 Species | *Pseudomonadota* |
| --- | --- | --- | --- |
| Document | Genomes | Species | Species |
| # of BWT Runs ($r$) | 19,088,436 | 865,907,683 | 25,243,943,020 |
| # of unique colors ($\vert C\vert $) | 338,796 | 1,137,989 | 2,477,411,592 |
| $r/\vert C\vert $ | 56.34 | 760.91 | 10.19 |

**Table 2.** Accuracy comparison with different threshold approaches. All the strategies perform similarly for multi-class classification of the reads.

| Repositioning Strategy | True thresholds | Mid-run thresholds | End of the run thresholds |
| --- | --- | --- | --- |
| Precision | 0.956 | 0.952 | 0.956 |
| Recall | 0.935 | 0.929 | 0.935 |
| F1 | 0.946 | 0.941 | 0.945 |

Note that, while *r/* |*C*| is greater than 10 in all the cases, it varies considerably across datasets. Two major factors drive this ratio; the total number of documents in the index, and the amount of redundancy between the documents.

As the total number of documents increases, the number of possible colors (the number of subsets of the documents) grows exponentially. In such cases, even a small fraction of possible colors can be very large. Even though the number of unique colors is bounded by the number of runs, it can get very close to *r* as the number of documents increases. That is the case with the *Pseudomonadota* dataset where there are 5, 326 distinct documents in the index. While the number of colors for this dataset is large, we observe that the color frequencies, the number of runs in which a color appears, have a highly skewed distribution. The cumulative percent of the runs covered by the 100, 000 most frequently occurring colors is shown in the Supplementary Figure 1.

Further, when there are greater similarities between the documents, more combinations of the documents have a chance to appear together, which increases the total number of unique colors. For the *Bacillus subtilis* dataset, many different groups of the documents share distinct similarities with each other because the documents are related by being genomes of the same species. Therefore, even though the number of documents in the *Bacillus subtilis* and the 48-species datasets are similar, the *r/*|*C*| for the 48-species data is much greater.

### 4.2 Evaluating the effect of alternative repositioning strategies

We first evaluated the effect of alternative repositioning strategies on the accuracy of Movi Color using simulated reads and an index built from the 48-species dataset. The read dataset includes positive reads, which originate from species represented in the index, and negative reads, which come from species that are not represented at any taxonomic level below the domain in the index. The positive reads are specifically simulated from strains that are not included in the index, although other strains from the same species are present.

This experiment is designed to reflect real-world scenarios where: (1) some genomes in the sample are completely novel and thus absent from the index, and (2) even for genomes that are represented, the exact strains generating the reads are typically not included in the index built from common reference databases.

We compared the accuracy of the classification with Movi Color when different repositioning strategies are used. We consider three different strategies: using the true thresholds, using the end of the run as the threshold (always repositioning to the up direction), or using the middle of the run as the threshold (mid-run thresholds). We observe that using alternative repositioning strategies does not impact the classification accuracy significantly. The result in Section 4.2 suggests that both precision and recall are impacted by less than one percent when an alternative strategy for repositioning is used. For exmample, the F1-score is between 0.941 to 0.946 for all cases. Thus, when the true thresholds are not available, we can switch to one of these alternative strategies for classification.

### 4.3 CAMI datasets with *Pseudomonadota* index

To benchmark the proposed method, we used the plant-associated metagenomic read dataset from the CAMI II challenge. We compared Movi Color with Kraken 2 (version 2.1.3) [30] and Metabuli (version v1-82ee9) [16].

We built the indexes of each tool (Movi, Kraken2, and Metabuli) over the *Pseudomonadota* reference dataset which consists of all complete genomes under the *Pseudomonadota* phylum in the NCBI database. In total, there were 5, 326 species, and 75, 166 distinct genomes.

Building Movi Color’s index requires the pre-processing of the reference sequences to compute the BWT. By default, Movi Color uses the “pfp-thresholds” software. This software implements the prefix-free parsing (PFP) algorithm [8], which is particularly efficient for building the BWTof a highly repetitive text such as a pangenome. “pfp-thresholds” also integrates Rossi et al’s [28] approach for computing thresholds required for repositioning. However, we were unable to run PFP on this dataset due to the high memory requirements imposed by the dataset’s diversity. Instead, we switch to an alternative method, grlBWT [12], which is also well-suited for building BWT of repetitive text. Unlike PFP, grlBWT makes use of disk space during construction to significantly reduce memory usage. A drawback of grlBWT is that it does not produce the thresholds data structure required for repositioning. As a result, Movi Color employs an alternative repositioning strategy (mid-run thresholds) when this construction method is used.

Since the indexes only include *Pseudomonadota* genomes (with *N* =), 927, 205 (out of 1, 691, 142) reads originate from a genome outside of the index. Furthermore, for evaluating the accuracy of the classification at each level, we only considered the reads that have a true label in the same or lower taxonomy level. For example, if the CAMI’s truth only assigns a taxonomy label to a read at the genus level, that read is not considered for evaluating the accuracy of the classification at the species level. Therefore, the set of reads being classified at each taxonomy level varies in their sizes ranging from 1,158,008 at the species level to 1,640,363 at the class level.

We performed the multi-class classification for the CAMI dataset. The results are reported in **Figure 3**. At the species rank, Movi Color outperforms Kraken 2 with twice the recall (0.138 vs 0.064) and with 65% higher precision (0.331 vs 0.200) **(Figure 3.a)**. Our results shows that Movi Color achieves a higher F1 score than Kraken 2 and Metabuli across all taxonomic levels, including species, genus, family, order, and class **Figure 3.b**.

**Figure 3.**
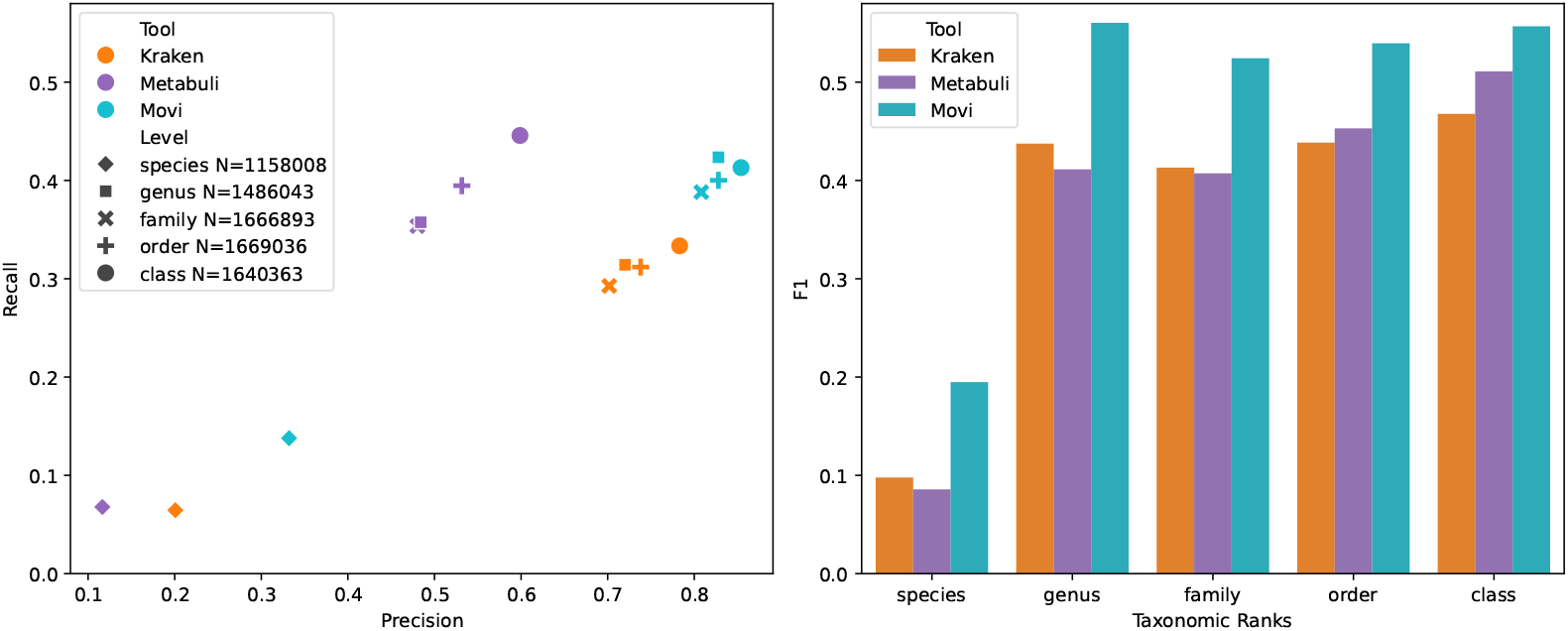
Classification accuracies of Movi Color compared to Metabuli and Kraken 2

We also evaluated the classification performance of the three tools on short-read sequencing data from CAMI. The results showed that Metabuli performed best at the species level, while Kraken was more effective at higher taxonomic levels. Movi demonstrated consistent performance across all levels (Supplementary Figure 2).

### 4.4 Evaluating runtime and memory usage

Here we compare the performance Movi Color with Kraken 2 and Metabuli in terms of runtime and memory requirements. We queried the CAMI II reads on each tool with their index built on the *Pseudomonadota* dataset.

We observe that the memory requirement for Movi Color is significantly higher than other tools. That is expected because Movi Color uses the a full-text index while tools like Kraken 2 use minimizer schemes for building a space efficient index.

When dealing with large indexes, a significant portion of the runtime is spent loading the index from disk into memory. Kraken 2 and Movi Color report the specific time spent only on read processing explicitly, so we are able to exactly compare the query time of their indexes. The columns under “Read processing” in Table 3 corresponds to this value exactly for both Kraken 2 and Movi Color. The index loading time for Movi Color is 11.77× longer compared to Kraken2 which is due to its size being 12.9× greater. For Metabuli, we were not able to exclude the index loading time, so the time reported in the read processing column includes both read processing and the index loading time. However, since Metabuli’s index size is closer to that of Kraken 2 rather than Movi Color, we expect its loading time to also be more comparable to Kraken 2.

**Table 3.**
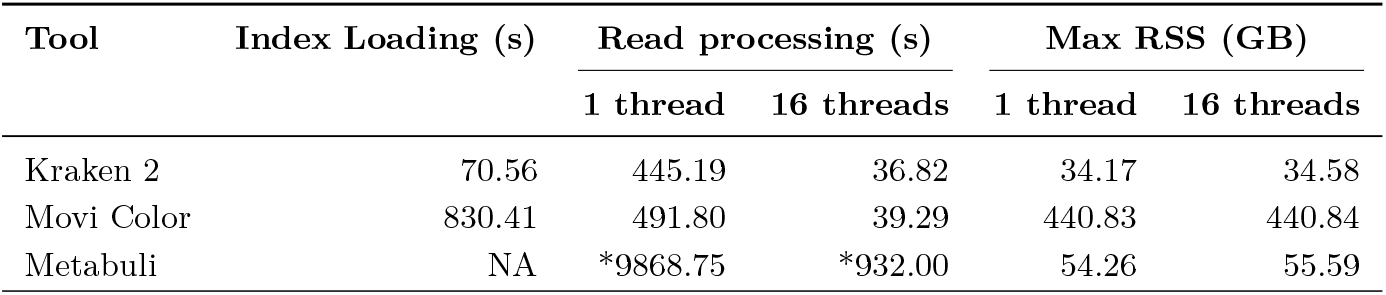
Runtime and maximum memory usage (Max RSS) for each classification tool using 1 and 16 threads with long CAMI II reads and the *Pseudomonadota* index. The size of Movi Color index can be broken into the colored move table (283 GB) and color table (157 GB). The indexes are built over all the complete genomes in the Phylum *Pseudomonadota* obtained from NCBI on November 9th, 2024 by Kraken 2. The index loading time for Kraken 2 is computed by decreasing the time reported for read processing from the total run time. Experiments were run on 3 GHz Intel Xeon Gold Cascade Lake 6248R CPU with 1.5TB DDR4 memory. All the timings are reported based on an average of 4 runs. *Metabuli does not report index loading time separately, so we reported the cumulative time including both index loading and classification.

| Tool | Index Loading (s) | Read processing (s) |  | Max RSS (GB) |  |
| --- | --- | --- | --- | --- | --- |
|  |  | 1 thread | 16 threads | 1 thread | 16 threads |
| Kraken 2 | 70.56 | 445.19 | 36.82 | 34.17 | 34.58 |
| Movi Color | 830.41 | 491.80 | 39.29 | 440.83 | 440.84 |
| Metabuli | NA | *9868.75 | *932.00 | 54.26 | 55.59 |

When we consider the read processing time, the time taken by Movi Color and Kraken2 are comparable. Movi Color’s runtime is only 10% slower than Kraken2 with 1 thread. The difference between the speed of Movi Color and Kraken2 goes down to 7% when 16 threads are used by both tools. Furthermore, the total runtime of Movi Color is 7.47 × faster than Metabuli using a single thread. Metabuli’s total runtime is more than 20× higher than the read processing time of Movi Color. We note that this performance is very impressive for a method based on a compressed full-text index and creates an opportunity for designing highly sensitive methods which are also very fast.

## 5 Discussion

We presented Movi Color, a new indexing strategy and software tool that uses the move structure to construct a highly efficient pangenome index. The index enables both multi-class classification queries and taxonomic classification queries. It exhibits superior classification accuracy compared to Kraken2 and Metabuli, at a speed that is comparable to Kraken2’s and much faster than Metabuli.

As noted in our results, there is still a substantial memory gap separating the Movi Color index from the Kraken 2 index for the same reference collection. This is undesirable, though it is also fully expected, since Movi Color is a full-text index while Kraken2’s index is tied to a particular minimizer scheme and k-mer length. Still, an important avenue for future work is to close the index-size gap between these tools. One promising strategy is to use *a priori* minimizer digestion [22] of the reference database and reads, similarly to SPUMONI2 [4].

Movi Color’s classification scheme mixes both the benefits and drawbacks of multi-class and taxonomic classification. The color information enables full “document listing” for matches, though only at a certain specified level of granularity. For instance, one project might use colors to describe documents at the species level whereas another might use color to describe documents at the genus level. When Movi Color is instead configured to report the best document match for the read, it behaves similarly to a multi-class classification tool like CLARK or SPUMONI2. When Movi Color is configured to report the lowest common ancestor of documents where matches occurred, it behaves similarly to a taxonomic classification tool like Kraken2. But unlike Kraken2, it is unable to report LCAs at “lower” (more specific) levels than the one used to define the documents. So, while we have removed the need for the user to select the correct k-mer length, there is still a need for the user to select which taxonomic level should be used for the colors. This also creates a trade-off space that the user must navigate: choosing a more specific level of the taxonomy can yield more specific and accurate classifications, but can also increase the size of the index since more distinct colors are needed to distinguish nodes at lower levels.

Furthermore, we note that there are many published schemes for compressing color information that could be adapted from the colored de-bruijn graph literature. In the future, we will investigate whether Movi Color could benefit from the additional strategies, like the meta colors [27] in Fulgor [13], or the efficient color representation using a spanning tree [5] in Mantis [26]. We will explore this in future work.

Finally, the compatibility scores computed by Movi Color for each read with respect to all the documents enable downstream applications such as metagenomics quantification and profiling. In the future, we plan to extend this framework by applying an expectation-maximization (EM) scheme that leverages these document-level scores from all the reads to estimate the relative abundances of the underlying documents.

## 6 Availability of data and materials

The CAMI read dataset is available at https://frl.publisso.de/data/frl:6425521/ plant_associated/long_read_pacbio/rhimgCAMI2_sample_0_reads.tar.gz. The scripts used for performing the experiments of the paper are available at https://github.com/mohsenzakeri/Movi-Color-experiments. Movi Color is available with the latest release of the Movi software https://github.com/mohsenzakeri/Movi/releases/tag/v1.2.0.

---

**Supplementary Figure 1.**
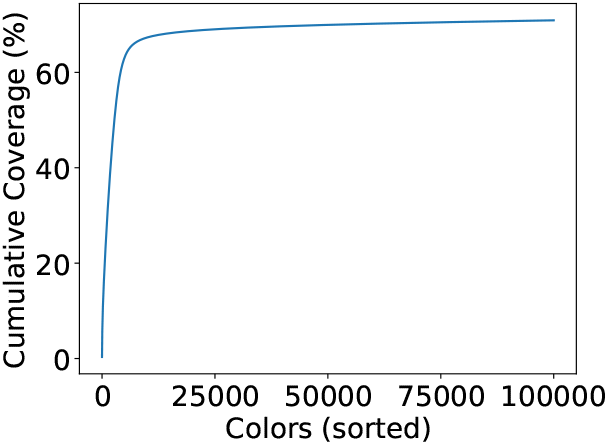
The cumulative coverage (% of runs) of the 10,000 most frequent colors (out of over 2.4 billion total colors) in index of *Pseudomonadota* reference genomes. The first 10, 000 most frequent colors cover 67.3% of the runs.

**Supplementary Figure 2.**
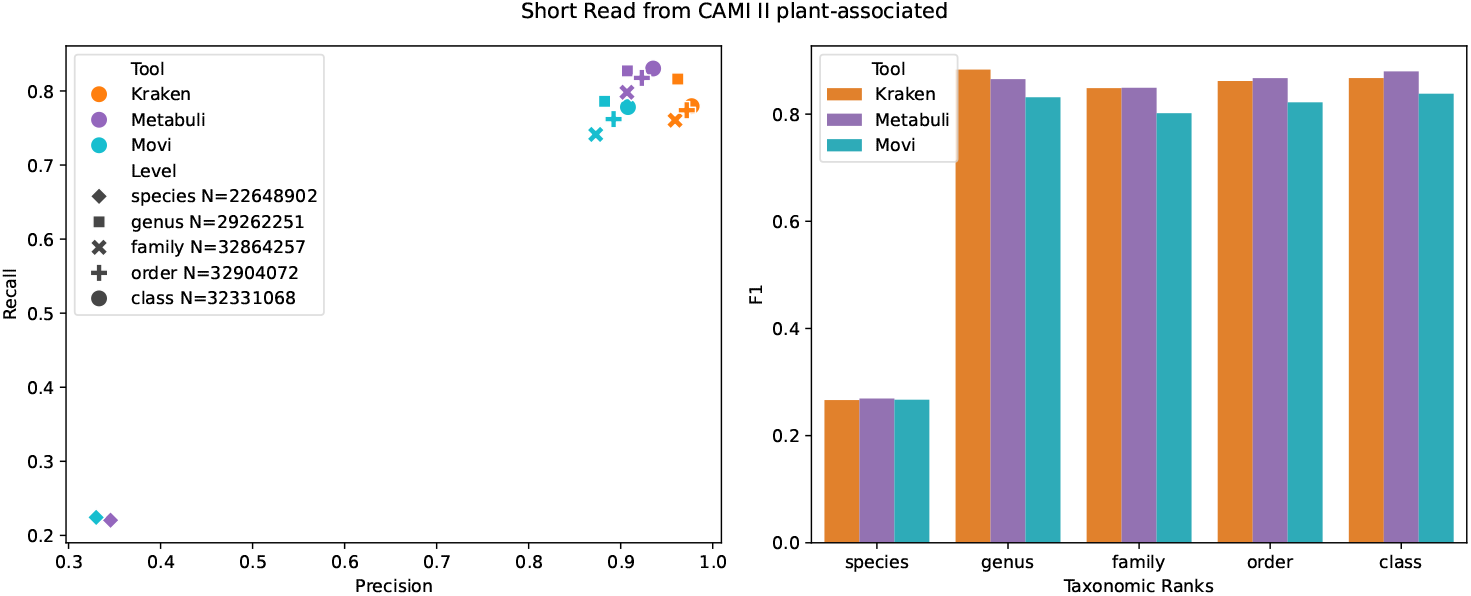
Classification accuracies of Movi Color compared to Metabuli and Kraken 2 on short reads.

2 For technical reasons, the move structure might partition some BWT runs[23]

